# Modelling the epidemic trend of the 2019 novel coronavirus outbreak in China

**DOI:** 10.1101/2020.01.23.916726

**Authors:** Mingwang Shen, Zhihang Peng, Yanni Xiao, Lei Zhang

## Abstract

We present a timely evaluation of the Chinese 2019-nCov epidemic in its initial phase, where 2019-nCov demonstrates comparable transmissibility but lower fatality rates than SARS and MERS. A quick diagnosis that leads to case isolation and integrated interventions will have a major impact on its future trend. Nevertheless, as China is facing its Spring Festival travel rush and the epidemic has spread beyond its borders, further investigation on its potential spatiotemporal transmission pattern and novel intervention strategies are warranted.

---

On 12^th^ December 2019, a pneumonia case of unknown etiology was reported in Wuhan City, Hubei Province, China, and on 31^st^ December 2019, the disease outbreak was reported to World Health Organization (WHO). After ruling out possible influenza and other coronaviruses by laboratory testing, the Chinese authorities isolated a new type of coronavirus (novel coronavirus, nCoV) on 7^th^ January 2020, which was then named 2019-nCoV by WHO on 12^th^ January [1]. As of 22^nd^ January, 571 confirmed cases (including 15 medical staff) and 17 deaths have been reported in China, and 6 cases confirmed overseas [2]. On 20^th^ January 2020. two local infections in the Chinese province of Guangdong with no direct visit to Wuhan were the first confirmed human-to-human transmission cases [3]. The asymptomatic incubation period (from infection to symptom onset) for individuals infected with 2019-nCov was 5-6 days (personal communication), and the virus infectiousness remained unknown. We estimated the basic and effective reproduction number of 2019-nCoV and predicted the epidemic peak time and size based on existing epidemiological data and a dynamic model.

Our mathematical model (details in Appendix) indicates that the national epidemic of 2019-nCov in China may lead to a total of 8042 (95%CI: 4199-11884) infections and 898 (368-1429) deaths (Figure 1a), corresponding to a fatality rate of 11.02% (9.26-12.78%). This is lower than the fatality rates of the Severe Acute Respiratory Syndrome (SARS) (14-15%) [4] and the Middle East Respiratory Syndrome (MERS) (34.4%) [5], suggesting that 2019-nCov may be a less virulent strain in the coronavirus family. Besides, experiences from fighting the previous coronaviruses may have also added to the rapid response to the epidemic by the Chinese government and the international society. However, we acknowledge that in the early phase of the epidemic, the death cases are likely under-reported as many infected cases have not progressed to the critical stage.

**Figure 1.**
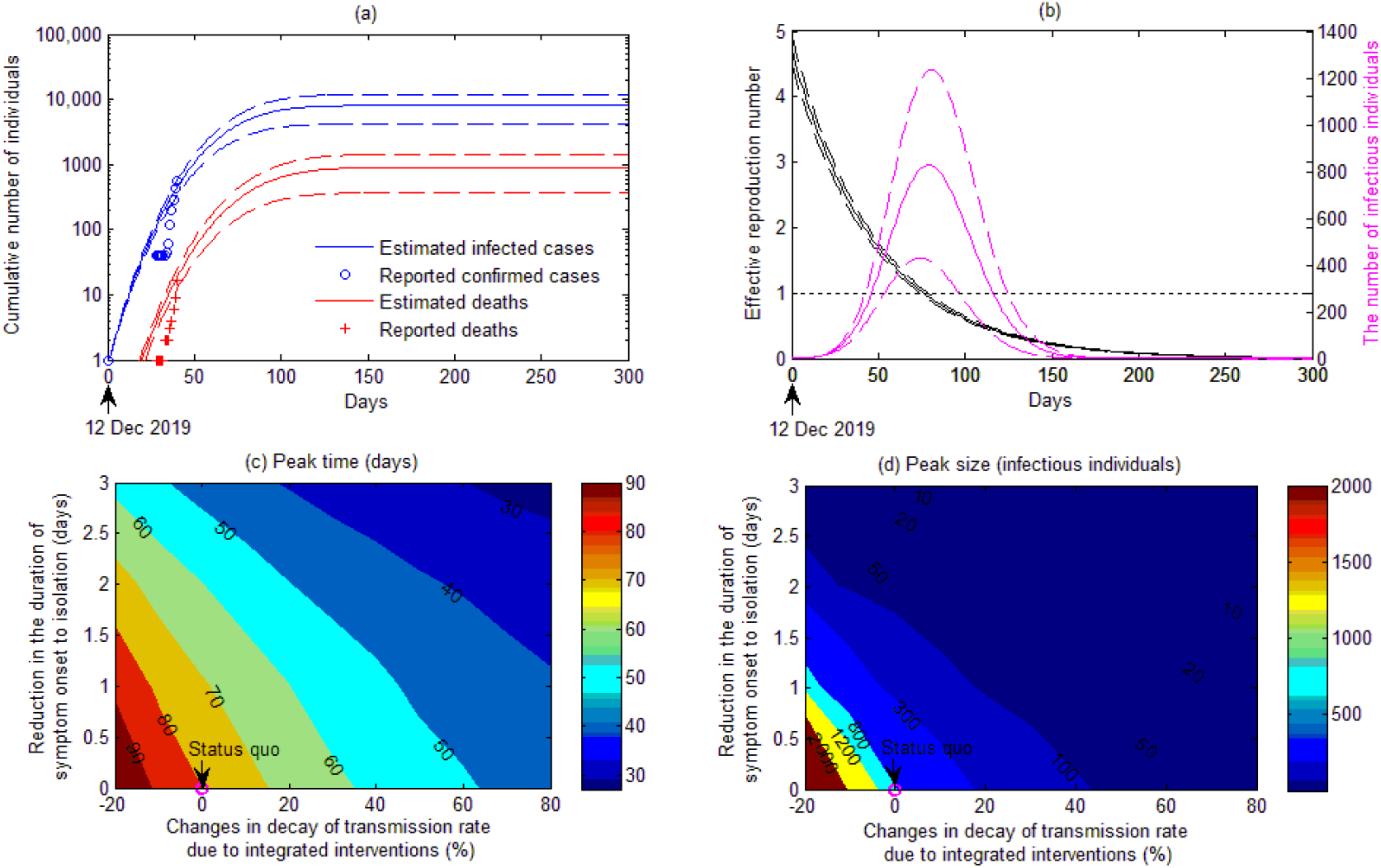
(a) Calibration of cumulative infected cases and death cases in China; the model is calibrated to reported data with 95% confidence intervals (CIs). (b) Estimated effective reproduction number and the number of infectious individuals over time (dashed lines denote 95% CIs and epidemic initiated on 12^th^ December 2019). Contour plot of peak time (c) and peak size (d) versus the time from symptom onset to isolation and changes in the decay of transmission rate due to integrated interventions.

The basic reproduction number (*R_0_*) of 2019-nCov, an indication of the initial transmissibility of the virus, was estimated to be 4.71 (4.50-4.92) when the epidemic started on 12^th^ December 2019, but its effective reproduction number (*R_e_*) has dropped to 2.08 (1.99-2.18) as of 22^th^ January 2020 (Figure 1b). If the declining trend continues with the assumption of no resurges of the epidemic, *R_e_* will drop below one within three months (77 [75-80] days) of the epidemic initiation, suggesting that the epidemic will gradually die off after this time. Compared with SARS and MERS, *R_0_* of 2019-nCov was similar to SARS (*R_0_*=4.91) in Beijing, China, in 2003 [6] and MERS in Jeddah (3.5-6.7) and Riyadh (2.0-2.8), Kingdom of Saudi Arabia, in 2014 [7].

Timely diagnosis for quarantine and integrated interventions are essential for curbing the epidemic. If the current intervention continues, the number of infected individuals is expected to peak in early March 2020 (80 days since initiation) with a peak population size of 827 (421-1232) infectious individuals in China. The current duration from symptom onset to isolation is about six days. Our model indicates that every one-day reduction in this duration would reduce the peak population size by 72-84% and the cumulative infected cases and deaths both by 68-80% (Figure 1c,d). Integrated interventions, such as the promotion of face mask use and reduction of travel, have been actively implemented. We estimate that every additional 10% decay in the transmission rate due to integrated interventions would reduce the peak population size by 20-47%, the cumulative infected cases and deaths both by 23-49% (Figure 1c,d). Facing the rapidly rising epidemic, the Chinese government has timely amended the *Law of the PRC on the Prevention and Treatment of Infectious Diseases* on 20^th^ January 2020 to include the 2019-nCov as a class-B infection but manage it as a class-A infection due to its severity [8]. Consequently, more than 30,000 PCR-fluorescence probing detection kit for 2019-nCoV RNA has been distributed to designated diagnosis centres in Wuhan [9]. The Chinese government has also taken an unprecedented action of locking down Wuhan and its nearby Huangguang city in a bid to minimise person-to-person contact on 23^rd^ January 2020 [10].

We present a timely evaluation of the Chinese 2019-nCov epidemic in its initial phase, where 2019-nCov demonstrates comparable transmissibility but lower fatality rates than SARS and MERS. A quick diagnosis that leads to quarantine and integrated interventions will have a major impact on its future trend. Nevertheless, as China is facing its ‘Spring Festival travel rush’ and the epidemic has spread beyond its borders, further investigation on its potential spatiotemporal transmission pattern and novel intervention strategies are warranted.

All authors declare that they have no competing interests.

## Acknowledgments

This work was supported by the National Natural Science Foundation of China (grant numbers: 8191101420(LZ), 11801435 (MS), 11631012 (YX), 81673275(ZP), 91846302(ZP)); Thousand Talents Plan Professorship for Young Scholars (grant number 3111500001); Xi’An Jiaotong University Young Talent Support Program; China Postdoctoral Science Foundation (grant number 2018M631134); the Fundamental Research Funds for the Central Universities (grant number xjh012019055); Natural Science Basic Research Program of Shaanxi Province (Grant number: 2019JQ-187); the National S&T Major Project Foundation of China (2018ZX10715002-004, 2018ZX10713001-001) and the Priority Academic Program Development of Jiangsu Higher Education Institutions (PAPD).

## Authors’ contributions

M.S., Z.P., Y.X., and L.Z. conceived and designed the study. M.S. analyzed the data, carried out the analysis and performed numerical simulations. M.S. wrote the first draft of the manuscript. M.S., Z.P., Y.X., and L.Z. contributed to writing the paper and agreed with manuscript results and conclusions.

