## Supplemental Appendix for "Modelling the epidemic trend of the 2019 novel coronavirus outbreak in China"

### Authors contribute equally.

This is a supplementary document describing mathematical modelling details presented in the main text and parameters estimation.

#### 1. Model formulation

We proposed a dynamic compartmental model to describe the transmission of 2019-nCov in China. The population was divided into five compartments: susceptible individuals (*S*), asymptomatic individuals during the incubation period (*E*), infectious individuals with symptoms (*I*), isolated individuals with treatment (*J*), and recovered individuals (*R*). The total population size was denoted as *N*, (*N*=*S*+*E*+*I*+*J*+*R*). Susceptible individuals became infected by being in contact with the infectious individuals and entered the latent compartment at the rate $\beta(t)SI/N$, where $\beta(t) = \beta e^{-m(t-\tau)}$, of which $\beta$ denoted the mean person-to-person transmission rate per day in the absence of control interventions, $\tau$denoted the time when the control interventions began, and *m* denoted the decay of transmission rate due to integrated interventions. Individuals in the incubation period progressed to the infectious compartment at a rate $k$, and infectious individuals were diagnosed and isolated at the rate $\alpha$. We assumed strict isolation that isolated individuals could not further infect others. Isolated individuals recovered at the rate $\gamma$or died due to the disease at the rate $\mu$. The model was described by the following system of ordinary differential equations:

$$\begin{cases} \frac{dS}{dt} = -\beta(t)\frac{SI}{N}, \\ \frac{dE}{dt} = \beta(t)\frac{SI}{N} - kE, \\ \frac{dI}{dt} = kE - \alpha I, \\ \frac{dJ}{dt} = \alpha I - (\gamma + \mu)J, \\ \frac{dR}{dt} = \gamma J. \end{cases} \quad (1)$$

The cumulative number of infected cases *C* and deaths *D* (*C* and *D* were not epidemiological states) were governed by the equations

$$\frac{dC}{dt} = kE, \frac{dD}{dt} = \mu J. \quad (2)$$

The basic ($R_0$) and effective ($R_e(t)$) reproduction numbers have been previously defined [1,2], they were $R_0 = \frac{\beta}{\alpha}$ and $R_e(t) = \frac{\beta(t)}{\alpha}\frac{S}{N}$.

#### 2. Data sources and parameter estimation

We obtained the number of cumulative confirmed cases and deaths from the Wuhan Municipal Health Commission [3] and the National Health Commission of the People's Republic of China [4] (Table S1). The first diagnosis was reported on 12th December 2019 and we used this date as the starting date for the epidemic ($t$=0). Data from this day until 22nd January 2020 were calibrated using Eq. (2) to estimate the unknown parameters. The mean incubation time for 2019-nCov was five days ($1/k$=5) and the mean 'time from symptoms onset to isolation' was six days ($1/\alpha$ =6) [5,6]. We used these two values as prior information for Markov chain Monte Carlo (MCMC) simulations [7]. The mean time from isolation to recovery is chosen as 6 days ($1/\gamma$ =6). The total population size in China was chosen as 1,400,050,000 as in 2019 [8]. We assumed the integrated interventions were implemented after the appearance of the first infected case, i.e., $\tau=0$. The initial values of the disease states were given as E(0)=0, I(0)=1, J(0)=0, R(0)=0, N(0)= 1,400,050,000.

We calibrated the model (Eq. (2)) to the cases and deaths data from 12th December 2019 to 22nd January 2020 by using nonlinear least squares method and thus we obtained the point estimate of the transmission rate $\beta$, the decay in transmission rate due to integrated interventions $m$, and the disease-induced death rate $\mu$. Then we used these estimated values as prior information in MCMC methods with a Metropolis-Hastings (M-H) algorithm [7] implemented by *Matlab* 2019. The algorithm was ran for 10,000 iterations with a burn-in (some iterations at the beginning of an MCMC run are threw away) of 5000 iterations, and we used the rest 5000 iterations to derive the mean value and 95% CI of parameters. We estimated the mean incubation time $1/k$ as 5.0265 (95%CI: 4.9337-5.1228) days, the mean 'time from symptoms onset to isolation' $1/\alpha$ as 6.1388 (5.9676-6.3200) days, the transmission rate $\beta$ as 0.7676 (0.7403-0.7949), the decay in transmission rate due to integrated interventions $m$ as 0.0204 (0.0202-0.0206), and the disease-induced death rate $\mu$ as 0.0207 (0.0170-0.0243).

Based on these estimated parameter values, we used the model (Eq. (1)-(2)) to forecast the epidemic trend (including cumulative cases and deaths, effective reproduction number, and the number of infectious individuals) over a 10-months period since the epidemic initiation (Figure 1a-b in the main text). We also explored how the reduction in the duration from symptoms onset to isolation ($1/\alpha$ reduces by 1, 2, 3 days) and changes in decay of transmission rate due to integrated interventions ($m$ varies between 0.8$m$ and 1.8$m$) will affect the peak time and peak size as shown in Figure 1c-d in the main text.

Table S1. Reported cumulative confirmed cases and deaths data in China [3,4].

| Date | Cases | Deaths |
|---|---|---|
| 2019-12-12 | 1 | 0 |
| 2020-1-10 | 41 | 1 |
| 2020-1-11 | 41 | 1 |
| 2020-1-12 | 41 | 1 |
| 2020-1-13 | 41 | 1 |
| 2020-1-14 | 41 | 1 |
| 2020-1-15 | 41 | 2 |
| 2020-1-16 | 45 | 2 |
| 2020-1-17 | 62 | 2 |
| 2020-1-18 | 121 | 3 |
| 2020-1-19 | 198 | 4 |
| 2020-1-20 | 291 | 6 |
| 2020-1-21 | 440 | 9 |
| 2020-1-22 | 571 | 17 |
